# Parkin knockout inhibits neuronal development via regulation of proteasomal degradation of p21

**DOI:** 10.1101/093294

**Authors:** Mi Hee Park, Hwa Jeong Lee, Hye Lim Lee, Dong Ju Son, Jung Hoon Ju, Byung Kook Hyun, Sung Hee Jung, Ju-Kyoung Song, Dong-Hoon Lee, Chul-Ju Hwang, Sang Bae Han, Sanghyeon Kim, Jin Tae Hong

## Abstract

PARK2 encodes for the E3 ubiquitin ligase parkin and iimplicates in the development of Parkinson’s disease (PD). Although the neuroprotective role of parkin is well known, the mechanism of parkin’s function in neural stem differentiation is not clear. Co-expressions network analysis showed that SNAP25 and BDNF were positively correlated with parkin, but negatively correlated with p21 in human patient brain. Therefore, we investigated a link between the ubiquitin E3 ligase parkin and proteasomal degradation of p21 for the control of neural stem cell differentiation. We discovered that p21 directly binds with parkin and is ubiquitinated by parkin resulting in the loss of cell differentiation ability. Tranfection of p21 shRNA in PARK2 KO mice significantly rescued the differentiation efficacy as well as SNAP25 and BDNF expression. We also defined the decreased p21 ubiquitination and differentiation ability were reversed after treatment with JNK inhibitor, SP600125 in PARK2 KO mice derived neural stem cells. Thus, the present study indicated that parkin knockout inhibits neural stem cell differentiation by JNK-dependent proteasomal degradation of p21.

**Summary statement:** The present study indicated that parkin knockout inhibits neural stem cell differentiation by JNK-dependent proteasomal degradation of p21.

## Introduction

Neural stem cells (NSCs) can be differentiated into several cell types such as neurons and glia (Faigle and Song, 2013; Galli et al., 2003). In the biology of NSCs, neurogenesis is affected by the ubiquitination system. Protein ubiquitination plays a key role in neural stem cell self-renewal and cellular differentiation (Shi et al., 2010; Vishwakarma et al., 2014). It provides an additional layer of stem cell regulation at the post-translational level and extends to multiple cellular processes, including the modulation of extracellular matrix composition, surface receptor trafficking and signaling, control of the cell cycle, transcription factor abundance, and deposition of epigenetic marks (Strikoudis et al., 2014). E3 ubiquitin ligase is a key regulator of NSC viability and differentiation. The absence of ubiquitin ligase in the brain caused severe impairment in stem cell differentiation and increased progenitor cell death. Impaired neurogenesis is found in severe neurodegenerative diseases such as Parkinson’s disease (PD) and Alzheimer’s disease (AD) (Choi et al., 2016; Le Grand et al., 2015; Oliveira et al., 2015).

Parkin is an E3 ubiquitin ligase and works in conjunction with E1 ubiquitin-activating enzymes and E2 ubiquitin-conjugating enzymes (Riley et al., 2013; Zhang et al., 2000). The process of E3 ligase ubiquitination targets damaged or unwanted proteins for degradation by the 26S proteasome (Goldberg, 2003; Śledź and Baumeister, 2016). Putative substrates of parkin include synphilin-1, CDC-rel1, cyclin E, p38 tRNA synthase, Pael-R, synaptotagmin XI, sp22 and parkin itself. Parkin has been implicated in a range of biological processes, including autophagy of damaged mitochondria (mitophagy), cell survival pathways, and vesicle trafficking (Imai et al.; Kawahara et al., 2008; Staropoli et al.). Several studies have suggested that parkin is also important for neurogenesis. A link between parkin and neurogenesis was first established in Drosophila, which displayed impairment in neuronal loss in an age-dependent manner (Romani-Aumedes et al., 2014). Neurogenesis is affected by Parkinson’s disease in both human patients and animal disease models and is regulated by dopamine and dopaminergic (DA) receptors, or by chronic neuroinflammation (Dauer and Przedborski, 2003; Hemmerle et al., 2012). The mutation of these genes is also related with neuronal cell differentiation. A recent study demonstrated that parkin mutations affect parkin solubility and impairs its E3 ligase activity, leading to a toxic accumulation of proteins within susceptible neurons that result in a slow but progressive neuronal degeneration and cell death (Romani-Aumedes et al., 2014). However, the clear mechanism of the function of parkin in neurogenesis from neural stem cell has not been fully studied yet.

p21^Waf1/Cip1^, an inhibitor of cyclin-dependent kinases, is required for proper cell-cycle progression (Abbas and Dutta, 2009b; Jung et al., 2010). p21 also mediates cellular senescence of stem cells (Gu et al., 2013). Inactivation of p21 disrupts quiescence in populations of adult hematopoietic and neural stem cells (Porlan et al., 2013). A recent study has indicated that p21 acts as a transcriptional repressor of Sox2 in NSCs, keeps Sox2 at physiological levels, and thereby prevents induced replicative stress and cell cycle withdrawal (Marqués-Torrejón et al., 2013a). Loss of p21 in NSCs results in increased levels of secreted BMP2 which induce premature terminal differentiation of multipotent NSCs into mature non-neurogenic astrocytes in an autocrine and/or paracrine manner (Porlan et al., 2013). In addition, p21 participates in a number of other specific protein–protein interactions that relate to cell-cycle control, apoptosis, and differentiation (Oh et al., 2002). This complexity of functions is well illustrated by the role of p21 in keratinocyte self-renewal and differentiation. Induction of p21 expression is also one of the earliest cell-cycle regulatory events that contribute to differentiation-associated growth arrest (Missero et al., 1995; Missero et al., 1996). p21 is required for the intrinsic commitment for differentiation of keratinocytes (Topley et al., 1999). At later stages of differentiation, the p21 protein is decreased by proteasome-mediated degradation, and this down-regulation is required for differentiation (Di Cunto et al., 1998). Sustained p21 expression under these conditions blocks terminal differentiation marker expression at the level of gene transcription (Devgan et al., 2006). Several reports suggest that p21 is tightly regulated at the transcriptional and posttranslational levels (Abbas and Dutta, 2009a; Jung et al., 2010). Several E3 ligases, such as SCF^skp2^, CRL4^cdt2^ and APC/C^cdc20^, target p21 for ubiquitin-mediated degradation at different stages of the cell cycle (Abbas et al., 2008; Amador et al., 2007; Wang et al., 2005). MKRN1 E3 ligase has also been shown to promote p21 ubiquitination and degradation (Lee et al., 2009). Although the ubiquitination of p21 has been reported to play a role in the regulation of stem cell differentiation, the interaction between a parkin E3 ligase and p21 in stem cell differentiation is unknown.

In the preliminary stage, we found that the p21 protein was highly accumulated in the neural stem cells of parkin mutant transgenic mice compared with that of non-transgenic mice. Therefore, we investigated the link between the ubiquitin E3 ligase parkin and proteasomal degradation of p21 for the control of neural stem cell differentiation in parkin knockout mice.

## Results

### Network data of parkin and neurogenesis-associated proteins

To explore novel pathogenic mechanisms of PD, we analyzed publicly available RNA-Seq data (29 PD, 44 controls) from the prefrontal cortex of individuals. The eigengene values of the control were significantly negatively associated with PD (Fig. S1A). We then conducted an unsupervised co-expression network analysis using Surrogate variable analysis (SVA) from all PD cases and controls in order to identify the co-expression modules associated with PD. Of the 17 co-expression modules that were generated with this combined data, 10 modules (R>0.3, p<0.05) were significantly associated with PD (S1B). These significant modules contained 2583 genes including genes related to synaptic transmission, ubiquitin-dependent protein catalytic process and neuronal differentiation (Fig. 1A). Among the 2583 genes, 40 genes including SNAP25 and BDNF are associated with neuronal differentiation. Moreover, PD related genes such as PARK2, SNCA and LRRK2 are associated with neuronal differentiation (Fig. 1B).

**Fig.1.**
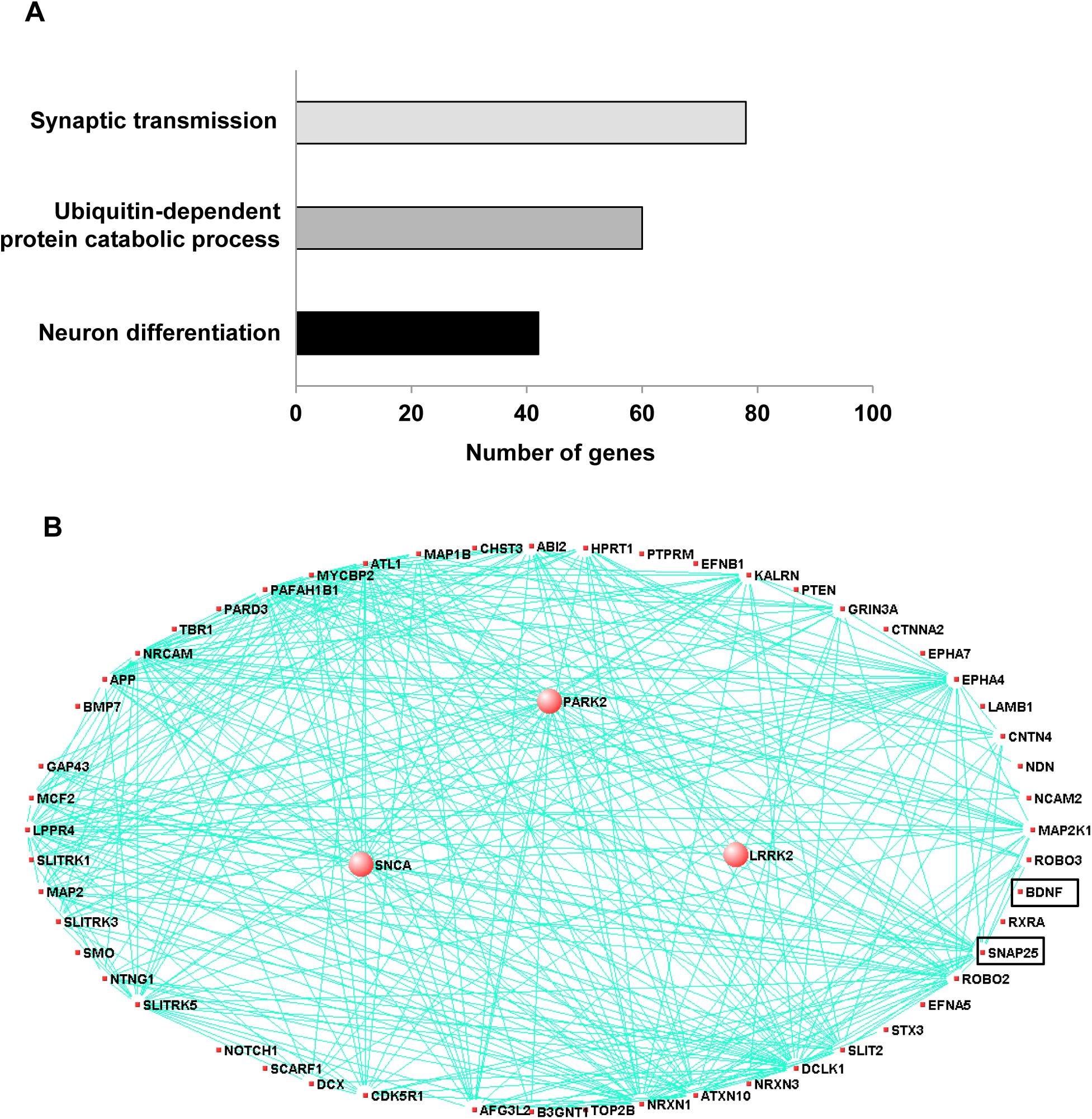
Co-expression module associated with Parkinson’s disease and related to neuron differentiation in prefrontal cortex. To explore novel pathogenic mechanisms of PD, we analyzed publicly available RNA-Seq data from the prefrontal cortex of individuals with 29 of PD and 44 of control. **A**, The network connections of three PD-associated genes and genes related neuron development and differentiation were visualized using VisANT. The three PD-associated genes are larger circles in the network. The eigengene values across samples in the PD_M17 module. Module PD_M17 was a large module containing 2583 genes and included three PD-associated genes such as PARK2 (parkin RBR E3 ubiquitin protein ligase), SNCA (synuclein alpha) and LRRK2 (leucine-rich repeat kinase 2). These genes are correlated with 54 of neuron differentiation related genes, especially SNAP-25 and BDNF. **B**, Major biological processes (Gene ontology) significantly enriched in the genes in the co-expression module.

### Effect of parkin on neurogenesis

In the network data, PD is related to the genes involved in neuronal differentiation and genes are related to the ubiquitin dependent protein. First, we investigated the effect of parkin on neurogenesis. To study the role of parkin on the spontaneous differentiation of neural stem cells, we cultured primary neural stem cells isolated from cortex of E15 days mouse embryos of non-tg or parkin knockout tg-mice. The cultured cells formed neurospheres at 5 days in vitro. We showed that neural stem cells from parkin knockout mice resisted spontaneous differentiation for a prolonged time, showed impaired differentiation, and were unsuccessful in forming a high-quality network of neural cells. We quantitatively analyzed the neurite of neural stem cells. We observed that neural stem cells from PARK2 knockout mice had significantly shorter length of primary and secondary neurites as well as a lower number of neurites per cell as compared to that of non-tg (Fig. 2A). In in vitro differentiation assay, we also found that TUBBIII-positive neuronal cells (Fig. 2B) and GFAP-positive astrocytes (Fig. 2C) were reduced in neural stem cells derived PARK2 knockout mice. PC12 has been previously used as an instructive model for studying the underlying mechanisms of neuronal differentiation in response to NGF (Teng et al., 1993). To check if the effect of parkin was similar in non-stem cells, PC12 cells were differentiated for 5 days upon stimulation with NGF (100 ng/ml) after the introduction of parkin shRNA. We showed that neurite-outgrowth and branching of PC12 cells were stimulated by the treatment of NGF, and this effect was inhibited by the treatment of parkin shRNA. In a quantified data, the average number of neurites per cell was much lower in parkin shRNA treated cells as compared to control cells (Fig. S2). Thus, parkin could be involved in neuronal differentiation both in the PC12 cell line as well as neural stem cells.

**Fig.2.**
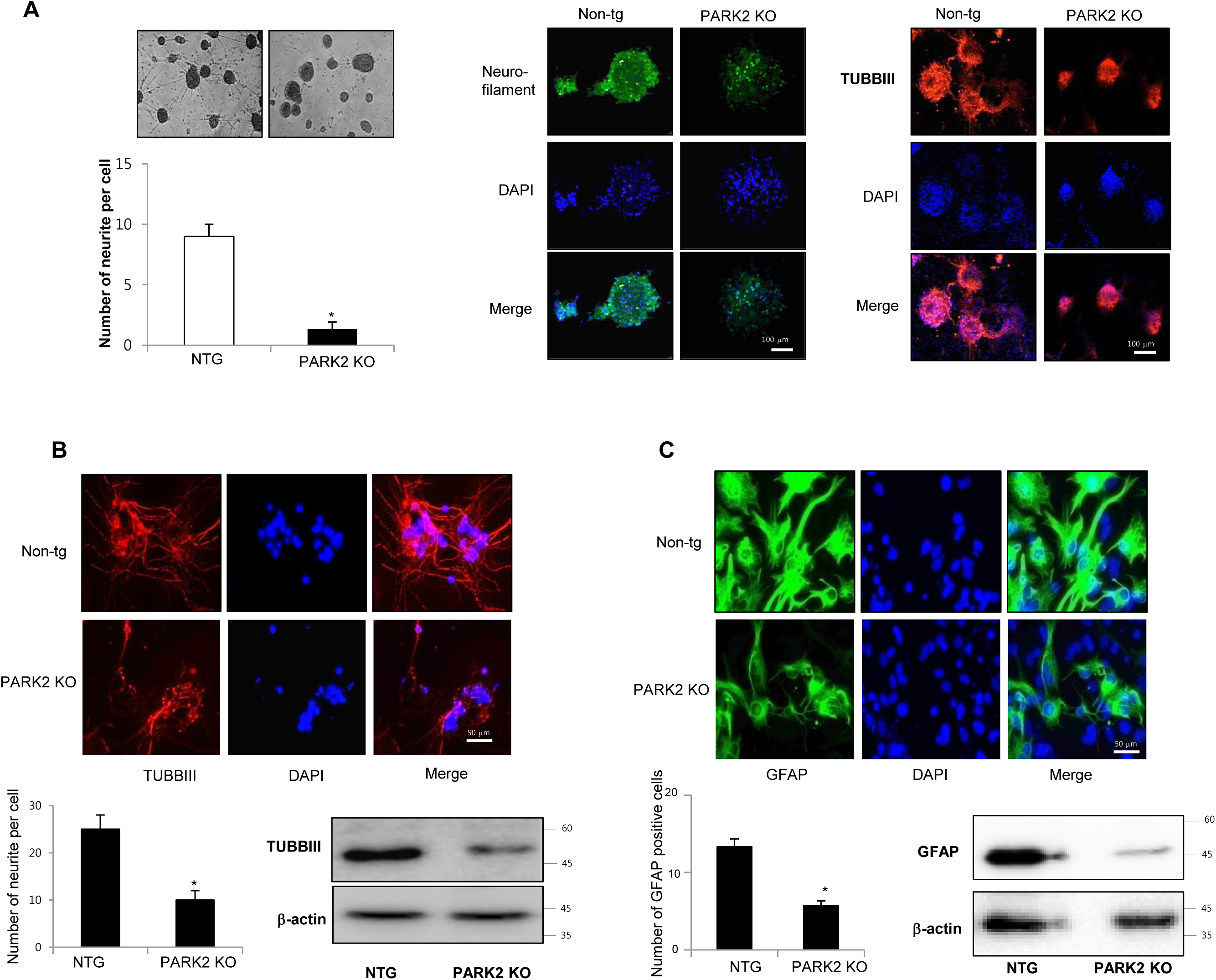
Effect of parkin on the differentiation of neural stem cells. **A,** Neural stem cells were isolated from embryonic day 15 forebrain germinal zones from parkin mutant or Non-tg mice. Neural stem cells were differentiated into astrocytes (**B)** and neuronal cells (**C**) in vitro as described in materials and methods. Western blot analysis confirmed the expression of TUBBIII and GFAP in terminally differentiated neurons and astrocytes. β-actin was internal control. Each band is representative for three experiments. The data are expressed as the mean ± SD of three experiments. *P < 0.05 compared with the non-tg derived neural stem cells.

### Involvement of p21 in the parkin regulated neuronal cell differentiation

Several recent studies showed that p21 is involved in stem cell neurogenesis. Moreover, in the analysis of human patients, the mRNA expression of SNAP25, a marker of synaptic formation, as well as BDNF, a marker of neurogenesis, was positively correlated with parkin, but negatively correlated with p21 (Fig. 3A). We found that among the cell growth regulatory genes such as p21, p53 and p27, p21 was significantly elevated in parkin KO mice derived neural stem cells (Fig. 3B). Thus, we examined whether p21 is associated with parkin-induced neurogenesis. We showed that the expression of SNAP25 and BDNF was downregulated in PARK2 KO mice-derived neural stem cells, but upregulated after treatment of p21 shRNA (Fig. 3B). We also found that the expression SNAP25 and BDNF was reversed by transfection of p21 shRNA in neural stem cells of PARK2 KO mice (Fig. 3C). This study indicated that parkin regulates neural stem cell differentiation through the regulation of p21. To understand the contribution of p21 on the parkin related differentiation of neural stem cells into the specific cell types, we determined specific neuronal cell differentiation. We found that the differentiated neuron and glial cells were much lower in parkin KO mice. The impaired differentiated cells in neural stem cells of PARK2 KO mice were reversed by transfection of p21 shRNA (Fig. 3D). Next, to determine the relationship of p21 and parkin in the differentiation of neural stem cells, we isolated the neural stem cells from non-tg or PARK2 KO mice and trasnsfected with p21 shRNA, and differentiated into astrocyte and neuronal cells. Cotransfection of p21 shRNA and parkin shRNA significantly rescued parkin shRNA induced inhibition of neurogenesis into neuronal cells and astrocytes. To confirm this phenomenon, we conducted the differentiation in vivo mouse model. We stereotaxically injected the parkin shRNA transfected-, p21 shRNA transfected- or co-transfected with parkin shRNA and p21 shRNA neural stem cells into the dorsal horn of the SVZ of 10-wk-old ICR mice. After 2 weeks following the injection, the injected neural stem cells were differentiated to astrocyte and neuron cells in the SVZ region of the brain by immunofluorescence staining. We showed that GFAP positive cells that are astrocyte cell markers are decreased in the parkin shRNA transfected cell injected group, but rescued in parkin shRNA and p21 shRNA cotransfected group (Fig. 3D). From this result, we suggest that p21 is critical for parkin-induced neurogenesis. These data indicate that p21 could be associated with parkin-induced neuronal differentiation.

**Fig.3.**
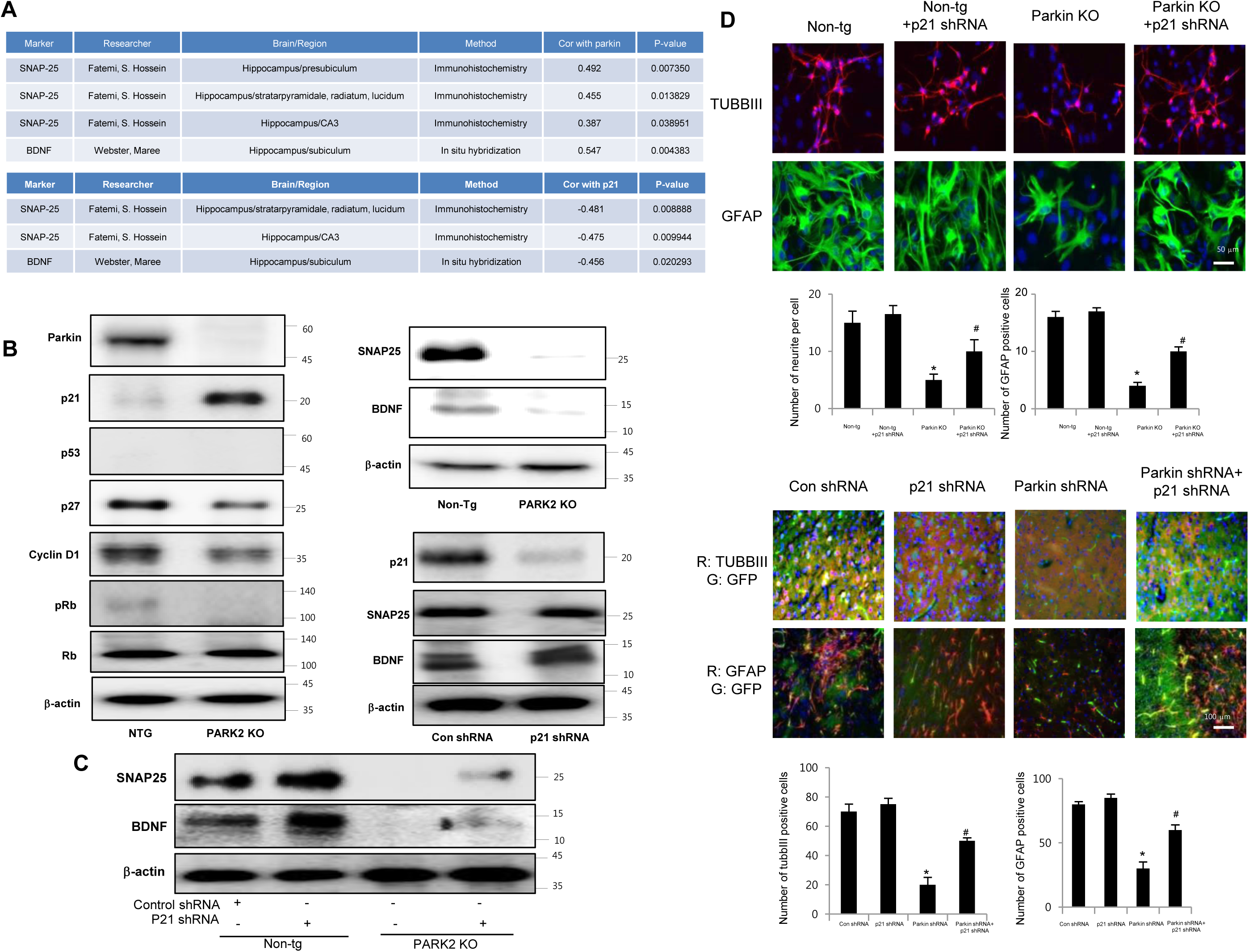
Effect of parkin and p21 on the expression of SNAP25 and BDNF. **A**, Correlation coefficients (R) and p-values between parkin or p21 and SNAP25 or BDNF. **B**, Neural stem cells isolated from Non-tg or PARK2 KO mice were harvested and western blotting was performed with p21, p53, p27, cyclinD1, pRb and Rb antibodies. Neural stem cells isolated from non-tg or PARK2 KO mice were differentiated for 5 days and then expression of SNAP25 or BDNF was visualized by Western blot analysis. **C**, Neural stem cells isolated from non-tg or PARK2 KO mice were transfected with p21 shRNA for 24hr, then differentiated, and then expression of SNAP25 or BDNF was visualized by Western blot analysis as described in the data. β-actin was internal control. Each band is representative for three experiments. **D**, Neural stem cells isolated from non-tg or PARK2 KO mice were transfected with p21 shRNA for 24hr, then differentiated into astrocytes and neuronal cells in vitro as described in materials and methods and then immunostained with GFAP or TUBBIII antibodies. Effect of p21 on the differentiation ability of parkin in vivo. Neural stem cells tranfected with parkin shRNA or p21 shRNA singly or together, and then injected in SVZ region of ICR mice. After 2 weeks, differentiated astrocyte or neuronal cells were stained with GFAP or TUBBIII. The data are expressed as the mean ± SD of three experiments. *P < 0.05 compared with the non-tg derived neural stem cells. ^#^P < 0.05 compared with PARK2 KO derived neural stem cells.

### Parkin ubiquitinates p21 in vivo and in vitro in an E3 ligase-substrate dependent manner

Parkin is involved in ubiquitination through their E3 ligase activity and p21 could be degraded by ubiquitin proteasome system. So, we questioned whether parkin is involved with the degradation of p21 in an ubiquitin proteasome system in the regulation of parkin-induced neurogenesis. We overexpressed Flag-tagged p21 and Myc-tagged parkin in HEK-293 cells and conducted immunoprecipitation studies to see whether parkin interacts with p21, thus degrades p21. We showed that parkin directly binds with p21 in the exogeneous HEK293 system (Fig. 4A upper panel). We also confirmed the interaction of p21 and parkin in primary neural stem cell cultures using coimmunoprecipitation experiments with parkin antibody (Fig. 4A lower panel). In the docking model, we showed that parkin binds with p21 at several sites including Leu187-Asn190, His279-Tyr285, Asn295-His302 and Tyr312-Cys323 (Fig.S3). These data indicate that parkin interacts with p21. To determine the functional significance of the interaction of p21 and parkin, we analyzed p21 levels in the presence of increasing amounts of parkin in HEK-293 cells. There was a dose-dependent decrease in steady-state levels of p21 as parkin levels increased (Fig. 4B upper panel). Furthermore, cycloheximide chase experiments indicated an accelerated decrease in steady-state levels of p21 when parkin was coexpressed with p21 in protein levels but not in RNA level (Fig. 4B lower panel). From this experiment, we expected that parkin influences on the p21 protein levels. Moreover, p21 is normally degraded by the ubiquitin proteasome system, and disruption of this pathway leads to increased p21 expression. We therefore set out to identify the parkin E3 ligase that ubiquitinates p21. To determine whether parkin directly ubiquitinates p21, we coexpressed p21 with hemagglutinin (HA)-ubiquitin and WT parkin and conducted ubiquitin affinity pull-down experiments. Overexpression of WT parkin resulted in an increase in ubiquitinated high-molecular-weight p21 bands (Fig. 4C left). We then conducted in vitro ubiquitination reactions, and found that p21 is directly ubiquitinated by parkin in the presence of the E1 ligase ubiquitin-activating enzyme E1 (UBE-1) and the E2 ligase ubiquitin-conjugating enzyme H7 (UBCHa5) (Fig. 4C right). We also determined the endogeneous ubiquitination in PARK2 KO derived neural stem cells. After the treatment of proteasome inhibitor MG132 (10 μM), we showed that ubiquitination was dramatically increased in PARK2 KO derived neural stem cells than that of Non-tg mice derived neural stem cells (Fig. 4D).

**Fig.4.**
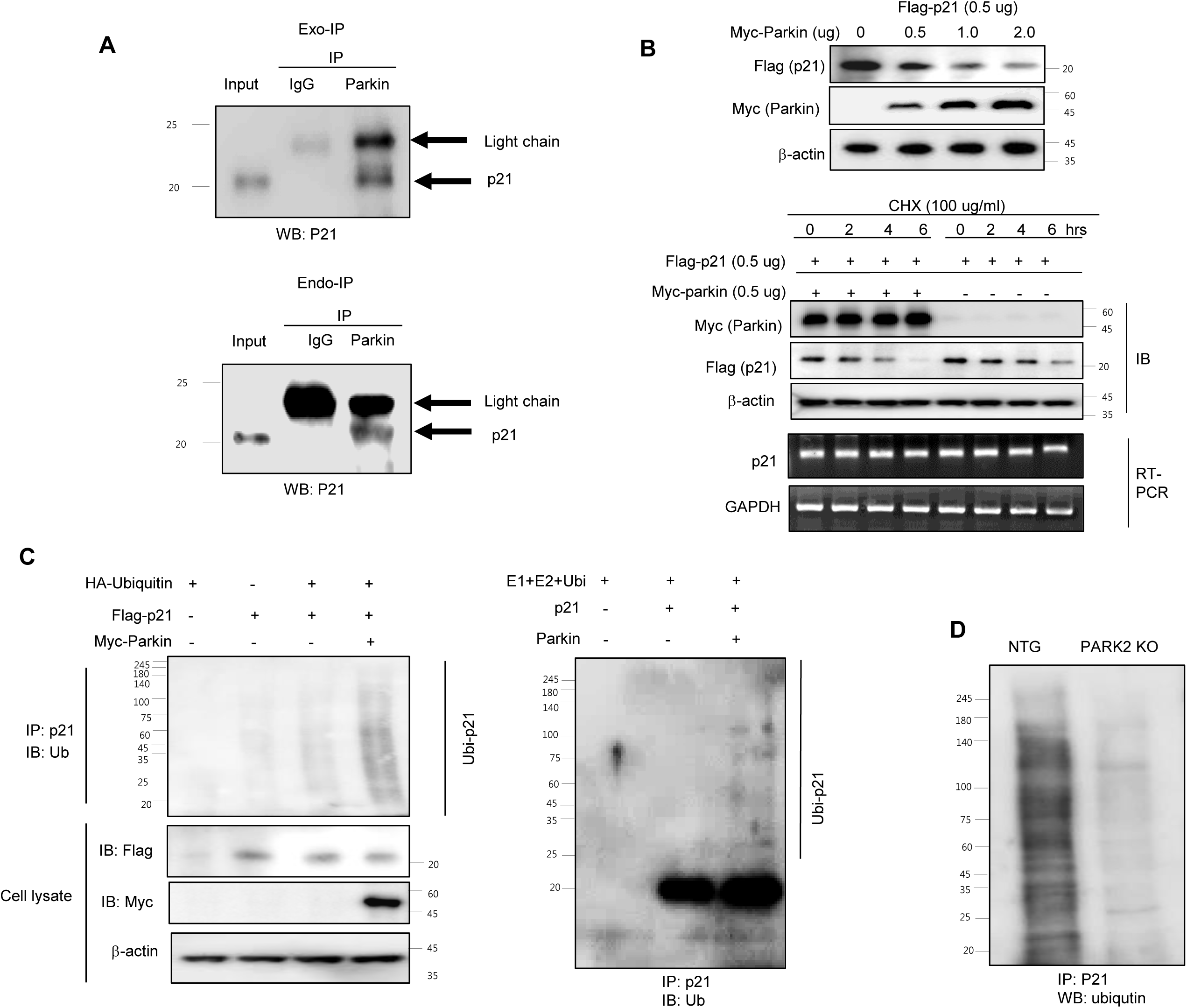
Parkin ubiquitinates p21 *in vivo* and *in vitro*. **A**, Myc-parkin and Flag-p21 were cotransfected into 239T cells. At 36 hours after transfection, cell lysates were harvested and was immunoprecipitated with anti-Myc or control IgG. Western blotting was performed (left panel). Parkin interacts with endogenous p21. Cell extracts of neural stem cells were immunoprecipitated with anti-Myc or control IgG. The immunoprecipitated complex was detected by Western blotting with anti-p21 (right panel). **B,** Parkin and ubiquitin proteasome system regulate p21 levels. Steady-state p21 levels were significantly decreased by coexpressing increasing amounts of parkin in HEK293 cells (upper panel). In the presence of cycloheximide, coexpression of parkin with p21 led to an accelerated decrease of p21 steady state levels compared with p21-alone transfected control (lower panel). For RT-PCR, total RNA were obtained from the transfected cells. The expression of p21 mRNA was determined by RT-PCR. GAPDH was internal control. **C**, 293T cells were cotransfected with constructs as indicated. At 48 hours after transfection, cells were treated with MG132 (10 μM) for 6hr, then harvested and immunoprecipitated with anti-p21. Ubiquitinated p21 was visualized by Western blot analysis using anti-ubiquitin (left panel). Parkin and Ubc5 mediate p21 ubiquitination. The ubiquitination reaction buffer, E1, UbcH5a, and parkin were incubated with p21 and polyubiquitinated p21 was detected by Western blotting (right panel). **D**, Endogeneous ubiquitination of p21 in non-tg or PARK2 KO mice. Neural stem cells isolated from non-tg or PARK2 KO mice were differentiated for 5 days, and then treated with MG132 (10 μM) for 6hr, then harvested and immunoprecipitated with anti-p21. Ubiquitinated p21 was visualized by Western blot analysis using anti-ubiquitin. Each band is representative for three experiments.

### Effect of parkin mutation on p21 levels and neural stem cell differentiation

To determine the effects of clinically relevant parkin mutations on p21 steady state levels and neuronal differentiation, we evaluated neuronal differentiation by two distinct types of genetic missense mutation; R275W that is in the RING1 domain and G430D that is in the RING2 domain of parkin both presumed to result in the loss of parkin function. We showed that p21 levels were significantly reduced by coexpression of WT parkin, which was blocked by the proteasome inhibitor cyclohexymide, but the expression of p21, along with familial-PD linked parkin mutants R275W or G430D, significantly recovered the steady-state p21 levels (Fig. 5A). Next, we coexpressed p21 with HA-ubiquitin and WT parkin or mutant parkin constructs (R275W, G430D) and conducted ubiquitin affinity pull-down experiments. Overexpression of WT parkin resulted in an increase in ubiquitinated high-molecular-weight p21 bands, however, coexpression of either parkin mutant resulted in a decrease in the levels of high-molecular-weight p21 species compared with WT parkin (Fig. 5B). Next, we checked the in vitro differentiation from neural stem cells by introducing parkin mutant plasmid. We showed that the differentiation of neural stem cells to astrocyte and neuron was inhibited by parkin mutation (Fig. 5C). These results further indicate the significance of p21 in parkin-induced neuronal differentiation (Fig.S4).

**Fig.5.**
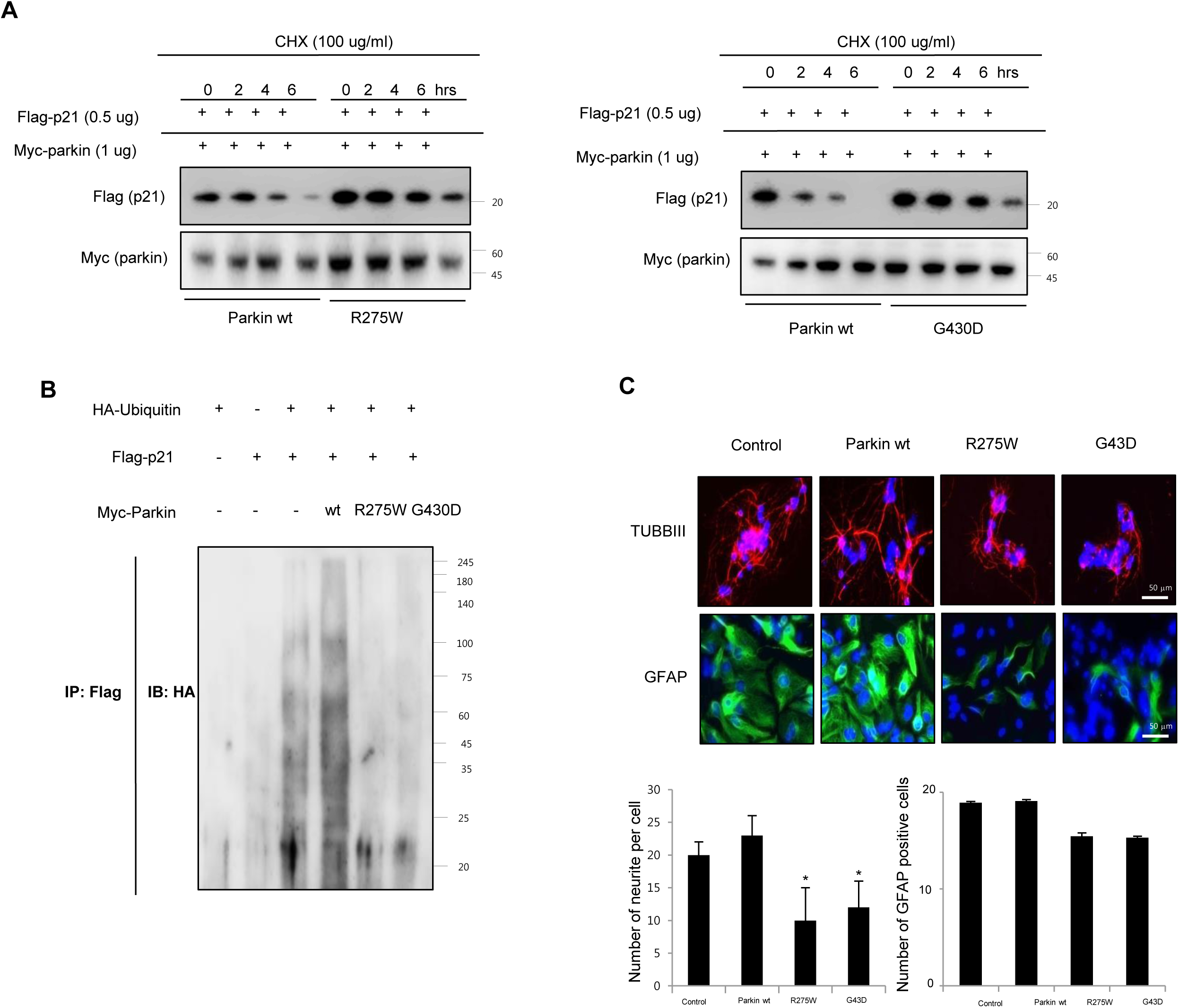
Effect of juvenile parkinson’s disease related parkin mutation on p21 levels and neural stem cell differentiation. **A,** WT parkin in HEK293 cells decreased the steady-state level of p21, but that was blocked by parkin mutants (R275W and G430D). **B**, Effect of parkin mutation on the ubiquitination of p21. 293 cells were cotransfected with constructs as indicated. At 48 hours after transfection, cells were harvested and then immunoprecipitated with anti-p21. Ubiquitinated p21 was visualized by Western blot analysis using anti-ubiquitin. Each band is representative for three experiments. **C**, Neural stem cells were transfected with parkin wt, parkin R275W or parkin G430D, then differentiate into astrocytes and neuronal cells in vitro as described in materials and methods and then immunostained with GFAP or TUBBIII antibodies. The data are expressed as the mean ± SD of three experiments. *P < 0.05 compared with the non-tg derived neural stem cells.

### pJNK is involved in parkin mediated p21 degradation

A recent study demonstrated that parkin inhibits the JNK signaling pathway in an E3 activity-dependent manner and suggested that loss of parkin function up-regulates the JNK signaling pathway in the dopaminergic neurons (40). In this study, we investigated whether JNK pathway is involved in parkin regulated p21 degradation. We showed that striatal pJNK levels are elevated in the neural stem cells derived from PARK2 KO mice as well as p21 elevation (Fig. 6A). Next, we defined whether pJNK is involved in parkin mediated p21 degradation. We showed that p21 levels are increased in the neural stem cells after transfection with parkin shRNA, but decreased by treatment with JNK inhibitor, SP600125 (0, 5, 10 μM) in a dose-dependent manner (Fig. 6B). We also checked whether JNK pathway was involved in parkin regulated p21 ubiquitination. We showed that p21 ubiquitination was decreased in parkin KO mice derived neural stem cells after a proteasome inhibitor, MG132 treatment, but reversed after the treatment with SP600125 (Fig. 6C). The differentiated astrocyte or neuronal cells are decreased from parkin knockdown neural stem cells, but reversed after the treatment with SP600125 in a dose-dependent manner (Fig. 6D). We also showed that striatal pJNK levels are elevated after MPTP treatment for the induction of Parkinson’s disease accompanied with elevated p21 levels (Fig. 6E). These data indicate the involvement of JNK pathway in the p21 pathway dependent neuronal differentiation function of parkin.

**Fig.6.**
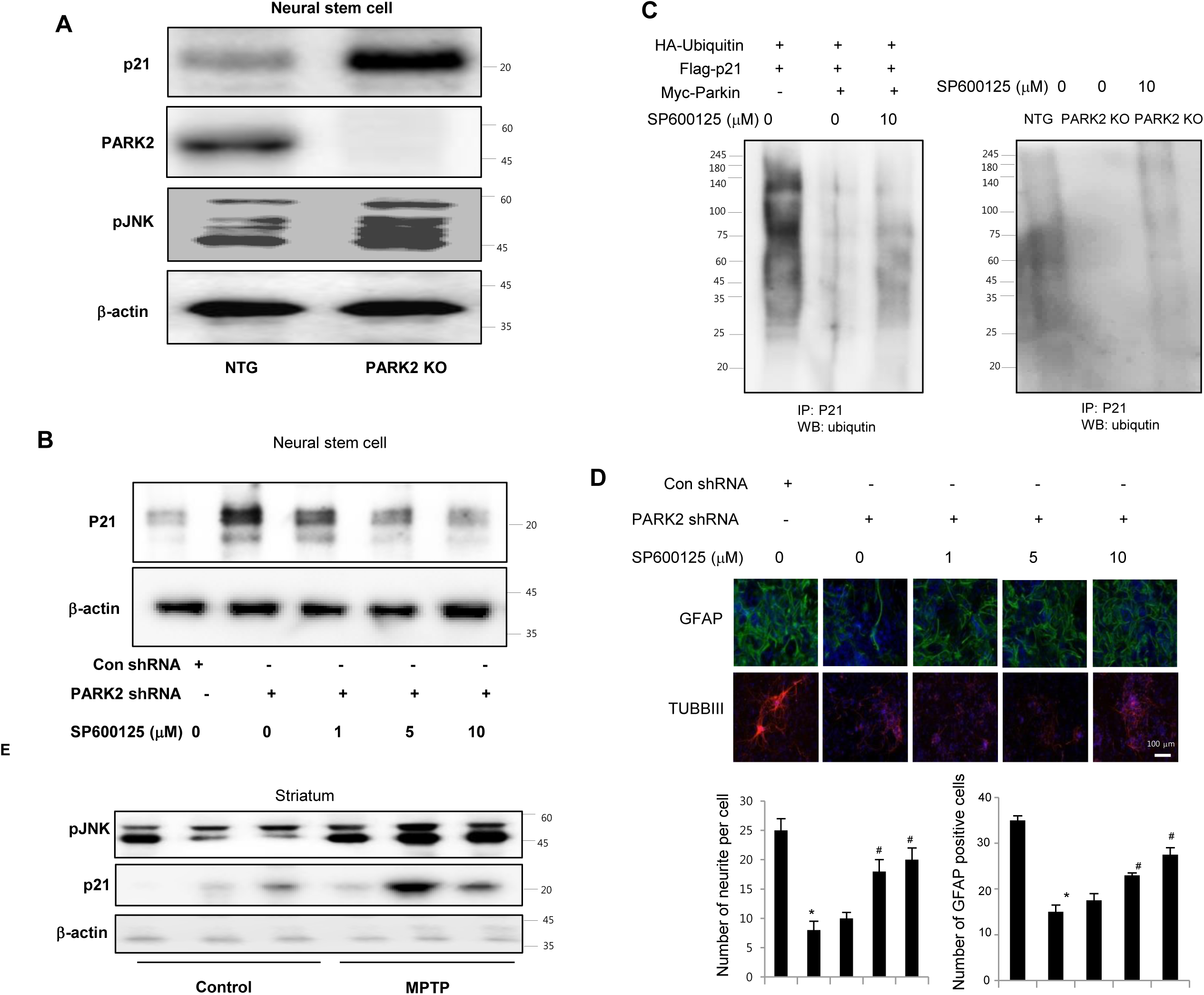
JNK is involved in parkin mediated p21 degradation. **A,** Expression of pJNK in neural stem cells of non-tg or PARK2 KO mice. **B**, Neural stem cells were transfected with parkin shRNA, and then treated with JNK inhibitor, SP600125 (0, 5, 10 μM) and then checked the p21 expression by western blotting. **C**, Neural stem cells were cotransfected with constructs as indicated. At 48 hours after transfection, cells were treated with SP600125 (10 μM) and MG132 (10 μM) for 6hr, then harvested and immunoprecipitated with anti-p21. Ubiquitinated p21 was visualized by Western blot analysis using anti-ubiquitin. **D**, Neural stem cells were transfected with parkin shRNA, and then treated with JNK inhibitor, SP600125 (0, 5, 10 μM) and then differentiated into astrocyte or neuronal cells. The data are expressed as the mean ± SD of three experiments. *P < 0.05 compared with the control shRNA treated groups. ^#^P < 0.05 compared with control of parkin shRNA treated groups. **E,** Striatal pJNK levels are elevated after MPTP treatment for the induction of parkinson’s disease. Immunoblot and quantification indicate that pJNK and p21 expression was increased in MPTP-treated mouse striatum compared with vehicle-treated controls. Each band is representative for three experiments.

## Discussion

Several reported studies have demonstrated that the default of neurogenesis could be associated with neurodegenerative diseases including Parkinson’s disease (Le Grand et al., 2015). A two-fold overexpression of α-synuclein that is involved in PD caused by a triplication of the SNCA gene is sufficient to impair the differentiation of neuronal progenitor cells (Oliveira *et al.,* 2015). PINK1 deficiency is also involved in PD decreasing brain development and neural stem cell differentiation (Choi *et al.,* 2016). They defined that the density of nestin- and beta3-tubulin-positive cells was reduced in the dentate gyrus (DG) of 3 patients with PD and in the DG of 5 patients with PD. These results suggested that Parkinson’s disease is associated with impaired neurogenesis.

In this study, we observed that differentiation of stem cells into neuronal cells and astrocytes were reduced in PARK2 KO mice derived neural stem cells, suggesting that parkin could be involved in neurogenesis. We also showed that neurite outgrowth of PC-12 cells was impaired in the PARK2 shRNA treated group. In addition, PD-associated PARK2 mutation including R275W and G430D also default differentiation of stem cells into neuronal cells and astrocytes. mRNA expression of neuronal differentiation associated genes such as SNAP25 as well as BDNF was also dramatically reduced in PARK2 KO mice derived neural stem cells. Moreover, co-expressions network analysis also showed that PD is related to neuron differentiation genes in the frontal cortex of PD patient brain. In particular, LRRK2 and SNCA are associated with parkin which are related to neurogenesis in PD. It has been reported that PD-associated LRRK2 mutation R1441G inhibits neuronal differentiation of neural stem cells (Bahnassawy et al., 2013). Elevated α-synuclein caused by SNCA is also involved in PD impairs neuronal differentiation. Therefore, these data indicated that PARK2 could be associated with other genes in the regulation of neural differentiation, and thus PARK2 knockout could inhibit neurogenesis. However, how PARK2 influence to neuronal differentiation is not yet understood.

p21 is known to be related with stem cell neurogenesis. A recent study demonstrated that cdkn1a-null cells induce the differentiation of neural stem cells, but reintroduction of p21 in neural stem cells reduced the terminal differentiation of astrocytes to wild type levels (Porlan et al., 2013). Sox2, an important factor for stem cell self-renewal and neurogenesis, is negatively related with p21 level (Marqués-Torrejón et al., 2013b). Additionally, elevated levels of BMP resulting from the lack of p21 induce terminal differentiation of neural stem cells into astrocytes (Porlan et al., 2013). These results indicated that p21 has an important role for stem cell selfrenewal and neurogenesis. RNA seq analysis showed that p21 is conversely correlated with neuronal differentiation marker SNAP25 and BDNF as well as parkin in PD patient brains. We demonstrated the link between the parkin and p21 for the control of neural stem cell neurogenesis. We showed that parkin directly binds with p21, ubiquitinates p21, and finally degrades p21 protein levels. Parkin shRNA-induced decreases differentiation ability of neural stem cells was rescued by cotransfection with p21 shRNA. Recent evidence verified that parkin mutations affect parkin solubility and impaired its E3 ligase activity, leading to a toxic accumulation of proteins resulting in a progressive neuronal degeneration and cell death (Romaní-Aumedes et al., 2014). It is also defined that that RTP801/REDD1, a pro-apoptotic negative regulator of survival kinases mTOR and Akt, is one of the parkin substrates which contributes to neurodegeneration caused by the loss of parkin expression or activity. p21 also mediates cellular differentiation (Gu et al., 2013). Loss of p21 in NSCs results in increased levels of secreted BMP2, which induce premature terminal differentiation of multipotent NSCs into mature non-neurogenic astrocytes in an autocrine and/or paracrine manner (Porlan et al., 2013). In addition, p21 is required for the intrinsic commitment for differentiation of keratinocytes (Topley et al., 1999). At later stages of differentiation, the p21 protein is decreased by proteasome-mediated degradation, and this down-regulation is required for differentiation (Di Cunto et al., 1998). Sustained p21 expression under these conditions blocks terminal differentiation marker expression at the level of gene transcription (Devgan et al., 2006). Several reports suggest that p21 is tightly regulated at the transcriptional and posttranslational levels (Abbas and Dutta, 2009a; Jung et al., 2010). These results suggest that-maintaining the level of p21 is important in parkin induced neural stem cell differentiation.

Other mechanisms may also be involved in the differentiation ability of parkin. The MAPK pathway is involved in the differentiation of neural stem cells. Moreover, one previous study indicated that pJNK can be regulated by parkin. Another recent study also suggested that parkin inhibits the JNK signaling pathway in an E3 activity-dependent manner in Drosophila parkin mutants (Cha et al., 2005). Phosphorylated c-Jun through the JNK pathway is an important regulator of neuronal viability. It was previously demonstrated that the depletion of Fbw7 leads to cell death in the neural stem cells cell due to increases in phosphorylated c-Jun through the JNK pathway (Hoeck et al., 2010). In agreement with this result, we also showed that the pJNK level was dramatically increased in parkin KO mice. We also defined whether pJNK is involved in parkin mediated p21 degradation. We showed that p21 levels increased in the neural stem cells after transfection with parkin shRNA, but decreased with the treatment of the JNK inhibitor SP600125 in a dose-dependent manner. The differentiated cells also decreased from parkin knockdown neural stem cells, but reversed after the treatment with SP600125 in a dose-dependent manner. In addition, the differentiated astrocytes and neuronal cells also decreased after the treatment with the JNK inhibitor. A recent study demonstrated that JNK is involved in p21 degradation. It has been reported that p21 is dynamically associated with JNK1 in vivo and JNK1 activation is correlated with dissociation of the p21WAF1/JNK1 complex in human T-lymphocytes (Patel et al., 1998). Moreover, JNK1 also can mediate increased stability of p21 protein by inducing phosphorylation at S130 in vivo and in vitro (Kim et al., 2002), and can regulate p21 level by proteasome/ubiquitin mediated degradation of p21. Taken together, our data indicate that JNK pathway, dependent on the degradation of p21, could also be involved in the neurogenesis by parkin.

## Materials and methods

### RNA-Seq data from prefrontal cortex of Parkinson’s disease patients and matched controls

RNA-Seq data from the prefrontal cortex of a previous human post-mortem study were downloaded from the GEO database. The data set (GSE8719) consists of a total of 73 normalized RNA-Seq data from the prefrontal cortex of individuals with PD (n=29) and from controls (n=44).

### Gene co-expression network analysis

We constructed co-expression networks using RNA-Seq data for all genes expressed in the prefrontal cortex and combined the data from both PD and control groups as previously described. Briefly, the potential confounding effects on the RNA-Seq data were identified using Surrogate variable analysis (SVA) and then the effects were adjusted on the data using the linear regression as previously described. The standardized residuals from the linear regression were used to generate gene co-expression networks using WGCNA. Correlation analysis between co-expression modules and diagnosis were performed to identify modules that were associated with the disease. P-values less than 0.05 were considered significant. The network connections were visualized using VisANT.

### Functional annotation and enrichment map

DAVID (http://david.abcc.ncifcrf.gov/home.jsp) was used to identify the biological processes that were significantly enriched in the genes included in the co-expression modules. P-values less than 0.05 were considered significant.

### Animals and isolation of neural stem cells

The parkin mutant mice were purchased from the Jackson Laboratory (Bar Harbor, ME, USA). The genetic background of non-transgenic mice and parkin mutant mice is C57BL/6. The mice were housed and bred under specific pathogen-free conditions at the Laboratory Animal Research Center of Chungbuk National University, Korea (CBNUA-929-16-01). Neural stem cells were isolated from embryonic day 14 forebrain germinal zones from parkin mutant or C57BL/6 (Jackson Laboratories).

Bulk cultures were established and the medium included DMEM/F12, 10% FBS and 1% penicillin/streptomycin. After 24hr, the medium was changed with a Neurobasal medium containing 1% glutamate, B27 supplement, 100 U/ml penicillin and 100 μg/ml streptomycin.

### PC12 cell culture and neurite outgrowth

PC12 cells were obtained from the American Type Culture Collection (Manassas, VA, USA). PC12 cells were grown in RPMI1640 with 5% FBS, 10% HS, 100 U/ml penicillin and 100 μg/ml streptomycin at 37 °C in 5% CO2 humidified air.

### Neurite outgrowth assay

To study neurite outgrowth, the medium was changed to RPMI containing 1% HS, 100 ng/ml NGF, 100 U/ml penicillin and 100 μg/ml streptomycin.

### Differentiation of neural stem cells into motor neurons

For in vitro priming, neural stem cells were cultured in neurobasal medium plus B27 (Invitrogen), 0.1 mM 2-mercaptoethanol, 20 ng/ml b-fibroblast growth factor, 1 μg/ml laminin, 5 μg/ml heparin, 10 ng/ml neural growth factor (Invitrogen), 10 ng/ml sonic hedgehog (R&D Systems), 10 μM forskolin (Sigma) and 1 μM retinoic acid (Sigma) for 5 days. For the induction of astrocyte, neural stem cells were cultured in poly-L-lysine coated slides with DMEM medium plus N2 supplement and L-glutamate. All cultures were maintained in a humidified incubator at 37°C and 5% CO2 in air, and half of the growth medium was replenished every third day.

### Transfection

Control shRNA, parkin shRNA, and p21 shRNA were purchased from Santa Cruz Biotechnology. The viral particles were combined with 8 μg/mL of polybrane (Santa Cruz Biotechnology) and infected overnight into neural stem cells or PC12 cells. The cell culture medium was replaced with fresh complete growth medium and after 48 hours, and the cells were used for experiments.

### Immunofluorescence staining

Immunofluorescence staining was done as previously described (Park et al., 2014).

### Immunohistochemistry

Immunohistochemistry was done as previously described (Park et al., 2014).

### Western blot

Western blot analysis was done as previously described (Park et al., 2014).

### Immunoprecipitation

Neural stem cells were gently lysed for 1 hr on ice and then centrifuged at 14,000 rpm 4 °C for 15 min, and the supernatant was collected. The soluble lysates were incubated with anti-parkin antibody (Santa Cruz Biotechnology Inc) at 4 °C for o/n and then with Protein A/G bead (Santa Cruz Biotechnology Inc) for 4 h at 4°C and washed 3 times. Immune complexes were eluted by boiling at 95 °C for 5 min in SDS sample buffer followed by immunoblotting with anti-p21 (1:500) antibody.

### RT-PCR

For mRNA quantification, total RNA was extracted using the easy-BLURTM total RNA extraction kit (iNtRON Biotech, Daejeon, Korea). cDNA was synthesized using High Capacity cDNA Reverse Transcription Kits (Applied Biosystems, Foster city, CA) according to the manufacturer’s instructions. Briefly, 2 μg of total RNA was used for cDNA preparation. RT-PCR was performed using the specific primers for p21 (Forward: 5’-GAGGCCGGGATGAGTTGGGAGGAG-3’ and Reverse: 5’-CAGCCGGCGTTTGGAGTGGTAGAA-3’) and GAPDH (Forward: 5’-GAAGGTGAAGGTCGGAGTC-3’ and Reverse: 5’GAAGATGGTGATGGGATTTC 3’).

### Docking procedure for parkin with p21

Molecular docking studies were performed using Autodock VINA. Parkin and p21 were obtained from the X-ray crystal structure. Parkin and p21 were used in the docking experiments and conditioned using AutodockTools by adding all polar hydrogen atoms. The grid box was centered on the parkin and the size of the grid box was adjusted to include the whole monomer. Docking experiments were performed at various exhaustiveness values of the default, 24, and 48. After the best binding mode was chosen, another round of docking experiments were performed with the grid box re-centered at the binding site of the best ligand-binding mode with its grid box size of 30 x 30 x 30.

### Ubiquitination assay

For in vivo ubiquitination assay, HEK293 cells were transfected with HA-Ub, Myc-parkin and/or Flag-p21 for 48 h, and then incubated with 10 μM MG132 for 4 h. Whole-cell lysates were co-immunoprecipitated with anti-Flag, and then analyzed by western blotting with anti-HA antibody. In vitro ubiquitination assays were performed as previously described (1). Briefly, recombinant parkin protein along with p21 (1 μg) were incubated for 3 hours at 37°C in a reaction buffer (20 mmol/L HEPES; pH 7.4, 10 mmol/L MgCl2, 1 mmol/L dithiothreitol, 59 mmol/L ubiquitin, 50 nmol/L E1, 850 nmol/L of UbcH5A and 1 mmol/L ATP, 30). After incubation, protein mixtures were diluted in an immunoprecipitation assay buffer and supernatant fractions were precleared with protein A/G beads for 2 hours, and immunoprecipitated overnight with anti-p21, after which protein A/G beads were added for an additional 2 hours. Beads were centrifuged and washed four times with E1A buffer. Proteins were eluted in 6 × SDS sample buffer and subjected to immunoblotting.

### Neural stem cell SVZ injections

Parkin shRNA transfected neural stem cells, p21 shRNA transfected neural stem cells, co-transfected with parkin shRNA and p21 shRNA neural stem cells or EV control (30,000 particles in 1 mL, four animals per condition) were stereotaxically injected into the dorsal horn of the SVZ of 10-wk-old ICR mice using coordinates AP +0.5; ML +1.1; from bregma and DV −1.9 from the pial surface. Mice were sacrificed 2 weeks following injection, and the CNS tissue was fixed by transcardial perfusion with PBS followed by 4% paraformaldehyde. To count cells in the olfactory bulb, five different fields at 203 were used from each animal.

### Statistical analysis

The data was analyzed using the GraphPad Prism 4 ver. 4.03 software (GraphPad Software, La Jolla, CA). Data is presented as mean ± SD. When the *p* value in the ANOVA test indicated statistical significance, the differences were evaluated by the Dunnett’s test. The value of *p* < 0.05 was considered to be statistically significant.

## Acknowledgement

This work was supported by the National Research Foundation of Korea (NRF) grant funded by the Korea government (MSIP) (No. MRC, 2008-0062275).

## Conflict of interest

The authors declare that they have no conflict of interest.

## Author contributions

JTH and SHK contributed to the design and coordination of the study. MHP performed all experiments. JTH, MHP and SHK participated in the study design and prepared the manuscript. LHJ, LHL and SDJ helped with image analysis and microscopy. JJH, HBK, JSH and SJK, advised with the isolation of neural stem cells from transgenic mice. LDH, HCJ and HSB participated in the technical supports. All authors read and approved the final manuscript prior to submission.

## References

Abbas, T. and Dutta, A. (2009a). p21 in cancer: intricate networks and multiple activities. Nat. Rev. Cancer 9, 400–414.

Abbas, T., Sivaprasad, U., Terai, K., Amador, V., Pagano, M. and Dutta, A. (2008). PCNA-dependent regulation of p21 ubiquitylation and degradation via the CRL4(Cdt2) ubiquitin ligase complex. Genes Dev. 22, 2496–2506.

Amador, V., Ge, S., Santamaría, P. G., Guardavaccaro, D. and Pagano, M. (2007). APC/C(Cdc20) controls the ubiquitin-mediated degradation of p21 in prometaphase. Mol. Cell 27, 462–473.

Bahnassawy, L. a., Nicklas, S., Palm, T., Menzl, I., Birzele, F., Gillardon, F. and Schwamborn, J. C. (2013). The Parkinson's Disease-Associated LRRK2 Mutation R1441G Inhibits Neuronal Differentiation of Neural Stem Cells. Stem Cells and Development 22, 2487–2496.

Cha, G.-H., Kim, S., Park, J., Lee, E., Kim, M., Lee, S. B., Kim, J. M., Chung, J. and Cho, K. S. (2005). Parkin negatively regulates JNK pathway in the dopaminergic neurons of Drosophila. Proceedings of the National Academy of Sciences of the United States of America 102, 10345–10350.

Choi, I., Woo, J. H., Jou, I. and Joe, E.-h. (2016). PINK1 Deficiency Decreases Expression Levels of mir-326, mir-330, and mir-3099 during Brain Development and Neural Stem Cell Differentiation. Exp Neurobiol 25, 14–23.

Dauer, W. and Przedborski, S. (2003). Parkinson’s Disease: Mechanisms and Models. Neuron 39, 889–909.

Devgan, V., Nguyen, B.-C., Oh, H. and Dotto, G. P. (2006). p21WAF1/Cip1 Suppresses Keratinocyte Differentiation Independently of the Cell Cycle through Transcriptional Up-regulation of the IGF-I Gene. Journal of Biological Chemistry 281, 30463–30470.

Di Cunto, F., Topley, G., Calautti, E., Hsiao, J., Ong, L., Seth, P. K. and Dotto, G. P. (1998). Inhibitory Function of p21Cip1/WAF1 in Differentiation of Primary Mouse Keratinocytes Independent of Cell Cycle Control. Science 280, 1069–1072.

Faigle, R. and Song, H. (2013). Signaling mechanisms regulating adult neural stem cells and neurogenesis. Biochimica et biophysica acta 1830, 2435–2448.

Galli, R., Gritti, A., Bonfanti, L. and Vescovi, A. L. (2003). Neural Stem Cells: An Overview. Circulation Research 92, 598–608.

Goldberg, A. L. (2003). Protein degradation and protection against misfolded or damaged proteins. Nature 426, 895–899.

Gu, Z., Jiang, J., Tan, W., Xia, Y., Cao, H., Meng, Y., Da, Z., Liu, H. and Cheng, C. (2013). p53/p21 Pathway Involved in Mediating Cellular Senescence of Bone Marrow-Derived Mesenchymal Stem Cells from Systemic Lupus Erythematosus Patients. Clinical and Developmental Immunology 2013, 134243.

Hemmerle, A. M., Herman, J. P. and Seroogy, K. B. (2012). Stress, Depression and Parkinson’s Disease. Experimental Neurology 233, 79–86.

Hoeck, J. D., Jandke, A., Blake, S. M., Nye, E., Spencer-Dene, B., Brandner, S. and Behrens, A. (2010). Fbw7 controls neural stem cell differentiation and progenitor apoptosis via Notch and c-Jun. Nat Neurosci 13, 1365–1372.

Imai, Y., Soda, M., Hatakeyama, S., Akagi, T., Hashikawa, T., Nakayama, K.-I. and Takahashi, R. CHIP Is Associated with Parkin, a Gene Responsible for Familial Parkinson’s Disease, and Enhances Its Ubiquitin Ligase Activity. Molecular Cell 10, 55–67.

Jung, Y.-S., Qian, Y. and Chen, X. (2010). Examination of the expanding pathways for the regulation of p21 expression and activity. Cellular signalling 22, 1003–1012.

Kawahara, K., Hashimoto, M., Bar-On, P., Ho, G. J., Crews, L., Mizuno, H., Rockenstein, E., Imam, S. Z. and Masliah, E. (2008). α-Synuclein Aggregates Interfere with Parkin Solubility and Distribution: ROLE IN THE PATHOGENESIS OF PARKINSON DISEASE. Journal of Biological Chemistry 283, 6979–6987.

Kim, G.-Y., Mercer, S. E., Ewton, D. Z., Yan, Z., Jin, K. and Friedman, E. (2002). The Stress-activated Protein Kinases p38α and JNK1 Stabilize p21Cip1 by Phosphorylation. Journal of Biological Chemistry 277, 29792–29802.

Le Grand, J. N., Gonzalez-Cano, L., Pavlou, M. A. and Schwamborn, J. C. (2015). Neural stem cells in Parkinson’s disease: a role for neurogenesis defects in onset and progression. Cellular and Molecular Life Sciences 72, 773–797.

Lee, E.-W., Lee, M.-S., Camus, S., Ghim, J., Yang, M.-R., Oh, W., Ha, N.-C., Lane, D. P. and Song, J. (2009). Differential regulation of p53 and p21 by MKRN1 E3 ligase controls cell cycle arrest and apoptosis. The EMBO Journal 28, 2100–2113.

Marqués-Torrejón, M. Á., Porlan, E., Banito, A., Gómez-Ibarlucea, E., Fernández-Capetillo, Ó., Vidal, A., Gil, J., Torres, J. and Fariñas, I. (2013a). Cyclin-dependent kinase inhibitor p21 controls adult neural stem cell expansion by regulating Sox2 gene expression. Cell stem cell 12, 88–100.

Marqués-Torrejón, M. Á., Porlan, E., Banito, A., Gómez-Ibarlucea, E., Lopez-Contreras, Andrés J., Fernández-Capetillo, Ó., Vidal, A., Gil, J., Torres, J. and Fariñas, I. (2013b). Cyclin-Dependent Kinase Inhibitor p21 Controls Adult Neural Stem Cell Expansion by Regulating Sox2 Gene Expression. Cell Stem Cell 12, 88–100.

Missero, C., Calautti, E., Eckner, R., Chin, J., Tsai, L. H., Livingston, D. M. and Dotto, G. P. (1995). Involvement of the cell-cycle inhibitor Cip1/WAF1 and the E1A-associated p300 protein in terminal differentiation. Proceedings of the National Academy of Sciences of the United States of America 92, 5451–5455.

Missero, C., Di Cunto, F., Kiyokawa, H., Koff, A. and Dotto, G. P. (1996). The absence of p21Cip1/WAF1 alters keratinocyte growth and differentiation and promotes ras-tumor progression. Genes & Development 10, 3065–3075.

Oh, H., Mammucari, C., Nenci, A., Cabodi, S., Cohen, S. N. and Dotto, G. P. (2002). Negative regulation of cell growth and differentiation by TSG101 through association with p21(Cip1/WAF1). Proceedings of the National Academy of Sciences of the United States of America 99, 5430–5435.

Oliveira, L. M. A., Falomir-Lockhart, L. J., Botelho, M. G., Lin, K. H., Wales, P., Koch, J. C., Gerhardt, E., Taschenberger, H., Outeiro, T. F., Lingor, P., et al. (2015). Elevated α-synuclein caused by SNCA gene triplication impairs neuronal differentiation and maturation in Parkinson’s patient-derived induced pluripotent stem cells. Cell Death & Disease 6, 093294.

Park, Y. S., Kang, J.-W., Lee, D. H., Kim, M. S., Bak, Y., Yang, Y., Lee, H. G., Hong, J. and Yoon, D.-Y. (2014). Interleukin-32α downregulates the activity of the B-cell CLL/lymphoma 6 protein by inhibiting protein kinase Cε-dependent SUMO-2 modification. Oncotarget 5, 8765–8777.

Patel, R., Bartosch, B. and Blank, J. L. (1998). p21WAF1 is dynamically associated with JNK in human T-lymphocytes during cell cycle progression. Journal of Cell Science 111, 2247–2255.

Porlan, E., Morante-Redolat, J. M., Marques-Torrejon, M. A., Andreu-Agullo, C., Carneiro, C., Gomez-Ibarlucea, E., Soto, A., Vidal, A., Ferron, S. R. and Farinas, I. (2013). Transcriptional repression of Bmp2 by p21Waf1/Cip1 links quiescence to neural stem cell maintenance. Nat Neurosci 16, 1567–1575.

Riley, B. E., Lougheed, J. C., Callaway, K., Velasquez, M., Brecht, E., Nguyen, L., Shaler, T., Walker, D., Yang, Y., Regnstrom, K., et al. (2013). Structure and function of Parkin E3 ubiquitin ligase reveals aspects of RING and HECT ligases. Nat Commun 4.

Romaní-Aumedes, J., Canal, M., Martín-Flores, N., Sun, X., Pérez-Fernández, V., Wewering, S., Fernández-Santiago, R., Ezquerra, M., Pont-Sunyer, C., Lafuente, A., et al. (2014). Parkin loss of function contributes to RTP801 elevation and neurodegeneration in Parkinson’s disease. Cell Death & Disease 5, 093294.

Romani-Aumedes, J., Canal, M., Martin-Flores, N., Sun, X., Perez-Fernandez, V., Wewering, S., Fernandez-Santiago, R., Ezquerra, M., Pont-Sunyer, C., Lafuente, A., et al. (2014). Parkin loss of function contributes to RTP801 elevation and neurodegeneration in Parkinson/’s disease. Cell Death Dis 5, 093294.

Shi, Y., Zhao, X., Hsieh, J., Wichterle, H., Impey, S., Banerjee, S., Neveu, P. and Kosik, K. S. (2010). microRNA regulation of neural stem cells and neurogenesis. The Journal of neuroscience: the official journal of the Society for Neuroscience 30, 14931–14936.

Śledź, P. and Baumeister, W. (2016). Structure-Driven Developments of 26S Proteasome Inhibitors. Annual Review of Pharmacology and Toxicology 56, 191–209.

Staropoli, J. F., McDermott, C., Martinat, C., Schulman, B., Demireva, E. and Abeliovich, A. Parkin Is a Component of an SCF-like Ubiquitin Ligase Complex and Protects Postmitotic Neurons from Kainate Excitotoxicity. Neuron 37, 735–749.

Strikoudis, A., Guillamot, M. and Aifantis, I. (2014). Regulation of stem cell function by protein ubiquitylation. EMBO Reports 15, 365–382.

Teng, K. K., Georgieff, I. S., Aletta, J. M., Nunez, J., Shelanski, M. L. and Greene, L. A. (1993). Characterization of a PC12 cell sub-clone (PC12-C41) with enhanced neurite outgrowth capacity: implications for a modulatory role of high molecular weight tau in neuritogenesis. Journal of Cell Science 106, 611–626.

Topley, G. I., Okuyama, R., Gonzales, J. G., Conti, C. and Dotto, G. P. (1999). p21(WAF1/Cip1) functions as a suppressor of malignant skin tumor formation and a determinant of keratinocyte stem-cell potential. Proceedings of the National Academy of Sciences of the United States of America 96, 9089–9094.

Vishwakarma, S. K., Bardia, A., Tiwari, S. K., Paspala, S. A. B. and Khan, A. A. (2014). Current concept in neural regeneration research: NSCs isolation, characterization and transplantation in various neurodegenerative diseases and stroke: A review. Journal of Advanced Research 5, 277–294.

Wang, W., Nacusi, L., Sheaff, R. J. and Liu, X. (2005). Ubiquitination of p21Cip1/WAF1 by SCFSkp2: Substrate Requirement and Ubiquitination Site Selection. Biochemistry 44, 14553–14564.

Zhang, Y., Gao, J., Chung, K. K. K., Huang, H., Dawson, V. L. and Dawson, T. M. (2000). Parkin functions as an E2-dependent ubiquitin– protein ligase and promotes the degradation of the synaptic vesicle-associated protein, CDCrel-1. Proceedings of the National Academy of Sciences of the United States of America 97, 13354–13359.

